# BAG3 and SYNPO (synaptopodin) facilitate phospho-MAPT/Tau degradation via autophagy in neuronal processes

**DOI:** 10.1101/518597

**Authors:** Changyi Ji, Maoping Tang, Claudia Zeidler, Jörg Höhfeld, Gail VW Johnson

**Affiliations:** Department of Anesthesiology, University of Rochester, 601 Elmwood Ave, Box 604, Rochester, NY 14642, USA; Institute for Cell Biology, University of Bonn, Ulrich-Haberland-Str. 61a, D-53121 Bonn, Germany

**Keywords:** autophagosome, postsynaptic density, PPxY domain, SQSTM1/p62, synapse, WW domain

## Abstract

A major cellular catabolic pathway in neurons is macroautophagy/autophagy, through which misfolded or aggregation-prone proteins are sequestered into autophagosomes that fuse with lysosomes, and are subsequently degraded. MAPT (microtubule associated protein tau) is one of the protein clients of autophagy. Given that accumulation of hyperphosphorylated MAPT contributes to the pathogenesis of Alzheimer disease and other tauopathies, decreasing endogenous MAPT levels has been shown to be beneficial to neuronal health in models of these diseases. A previous study demonstrated that the HSPA/HSP70 co-chaperone BAG3 (BCL2 associated athanogene 3) facilitates endogenous MAPT clearance through autophagy. These findings prompted us to further investigate the mechanisms underlying BAG3-mediated autophagy in the degradation of endogenous MAPT. Here we demonstrate for the first time that BAG3 plays an important role in autophagic flux in the neuritic processes of mature neurons (20-24 days in vitro [DIV]) through interaction with the post-synaptic cytoskeleton protein SYNPO (synaptopodin). Loss of either BAG3 or SYNPO impeded the fusion of autophagosomes and lysosomes predominantly in the post-synaptic compartment. A block of autophagy leads to accumulation of the autophagic receptor protein SQSTM1/p62 (sequestosome 1) as well as MAPT phosphorylated at Ser262 (p-Ser262). Furthermore, p-Ser262 appears to accumulate in autophagosomes at post-synaptic densities. Overall these data provide evidence of a novel role for the co-chaperone BAG3 in synapses. In cooperation with SYNPO, it functions as part of a surveillance complex that facilitates the autophagic clearance of MAPT p-Ser262, and possibly other MAPT species at the post-synapse. This appears to be crucial for the maintenance of a healthy, functional synapse.

## Introduction

Autophagy is an essential component of the protein quality control machinery that mediates the degradation and recycling of damaged or dysfunctional cytosolic constitutes [1]. In this process, cargos such as long-lived proteins, lipids, carbohydrates or damaged cellular components are engulfed in double-membrane vesicles, which subsequently fuse with lysosomes to form autolysosomes for degradation [1, 2]. Previous studies have demonstrated that autophagic activity is essential for the survival of post-mitotic neural cells, such as neurons. In mouse models, loss of basal autophagy causes neurodegeneration, significant behavioral deficits and premature death [3, 4]. Further studies found basal autophagic biogenesis is constitutively active and differentially regulated in distinct subcellular compartments in neurons [5]. The molecular basis of presynaptic autophagy or autophagy in neuronal soma has been closely examined [5–11]. However, mechanisms of the initiation and maturation of autophagy in the postsynaptic regions have not been reported.

The autophagic process was initially perceived as “in bulk” [2]. However, emerging evidence clearly suggests that selective autophagy plays a significant role in the degradation of misfolded or aggregation-prone proteins [12, 13]. Selective autophagy is supported by chaperone and cochaperones, which cooperate with autophagy receptors to direct specific cargos into autophagic vesicles [13]. For example, HSPA/HSP70 and its co-chaperone BAG3 (BCL2 associated athanogene 3) cooperate with the autophagy receptor SQSTM1/p62 to mediate the degradation of aggregation-prone proteins such as mutant huntingtin and superoxide dismutase 1 [13, 14]. In young neurons, soluble MAPT and its phosphorylated forms are also substrates of BAG3-mediated autophagy [15]. However, it remains unclear how BAG3 coordinates the autophagic machinery to mediate MAPT degradation in neurons.

BAG3 contains multiple domains including a HSPA/HSP70 interaction (BAG) domain at its C terminus, a tryptophan-containing (WW) domain at its N terminus, 2 IPV domains which mediate the interaction with small heat shock proteins such as HSPB6 and HSPB8, and a proline-rich (PXXP) domain in the central region [16, 17]. One of the important functions of BAG3 is to regulate selective autophagy via specific interactions with its partner proteins [17]. Previous studies have shown that the WW domain of BAG3 is required for autophagy induction in glioma cells and muscle cells [18, 19]. In particular, the WW domain of BAG3 interacts with a proline-rich motif (PPPY) in SYNPO2/myopodin (synaptopodin 2), an actin cytoskeleton adaptor protein, which facilitates the formation of autophagosomes during mechanical stress in muscle cells [18].

Neurons express a close homolog of SYNPO2, SYNPO (synaptopodin) [20, 21]. SYNPO is a post-synaptic protein that has been implicated in the structural and functional plasticity of dendritic spines [22–24]. SYNPO also acts as an organizer of the actin cytoskeleton at spines [25], which controls calcium release from local stores [26]. Of note, SYNPO contains 2 proline-rich motifs, one of which (PPSY) is a preferable binding motif of the BAG3 WW domain [18], raising the possibility that SYNPO interacts with BAG3 in neurons, and may cooperate with the co-chaperone during selective autophagy.

In this paper, we demonstrate for the first time that BAG3 and SYNPO interact in mature cortical neurons. The presence of the autophagy marker MAP1LC3B/LC3B as well as the autophagy receptor SQSTM1 in the BAG3-SYNPO complex indicated a role for the BAG3 and SYNPO interaction in the autophagic process. Interestingly, the interaction between BAG3 and SYNPO did not significantly affect the formation of autophagosomes, as previously shown for the BAG3-SYNPO2 interaction [18], but rather the fusion between autophagosomes and lysosomes. Knockdown of BAG3 or SYNPO affected autophagy only in neuronal processes, and in particular at the post-synapse, not in neuronal soma, which corroborated the compartment-specific regulation of autophagy in neurons. Furthermore, loss of SYNPO or BAG3 lead to accumulation of MAPT phosphorylated at Ser262 (p-Ser262) at postsynaptic densities; the level of other MAPT species (e.g. total MAPT, p-Thr231 or p-Ser396/404) was not altered. These observations suggest a dendritic specific mechanism of MAPT clearance via BAG3 and SYNPO assisted autophagy.

## Results

### BAG3 interacts with SYNPO in primary cortical neurons

The WW domain of BAG3 interacts with proline-rich PPxY motifs in partner proteins [18, 27] (Figure 1A). We recently performed a peptide array screen, which monitored binding of the purified WW domain to 2,296 PPxY-containing peptides derived from the human proteome [18]. The screen led to the identification of the cytoskeleton-associated protein SYNPO2 as a BAG3 binding partner [18]. In muscle cells, SYNPO2 links the BAG3 chaperone complex to VPS18-containing membrane fusion machinery, which facilitates autophagosome formation at sites of cytoskeleton damage [18]. The same screen also revealed binding of the BAG3 WW domain to a 12-mer peptide containing the PPSY motif of the SNYPO2-related protein SYNPO (amino acids [aa] 333-344) (Figure 1A). A second PPxY motif of SYNPO (PPTY, aa 318-312) was not among the 72 top hits of the peptide array screen, reflecting the fact that BAG3 preferentially binds PPPY and PPSY motifs [18].

**Figure 1.**
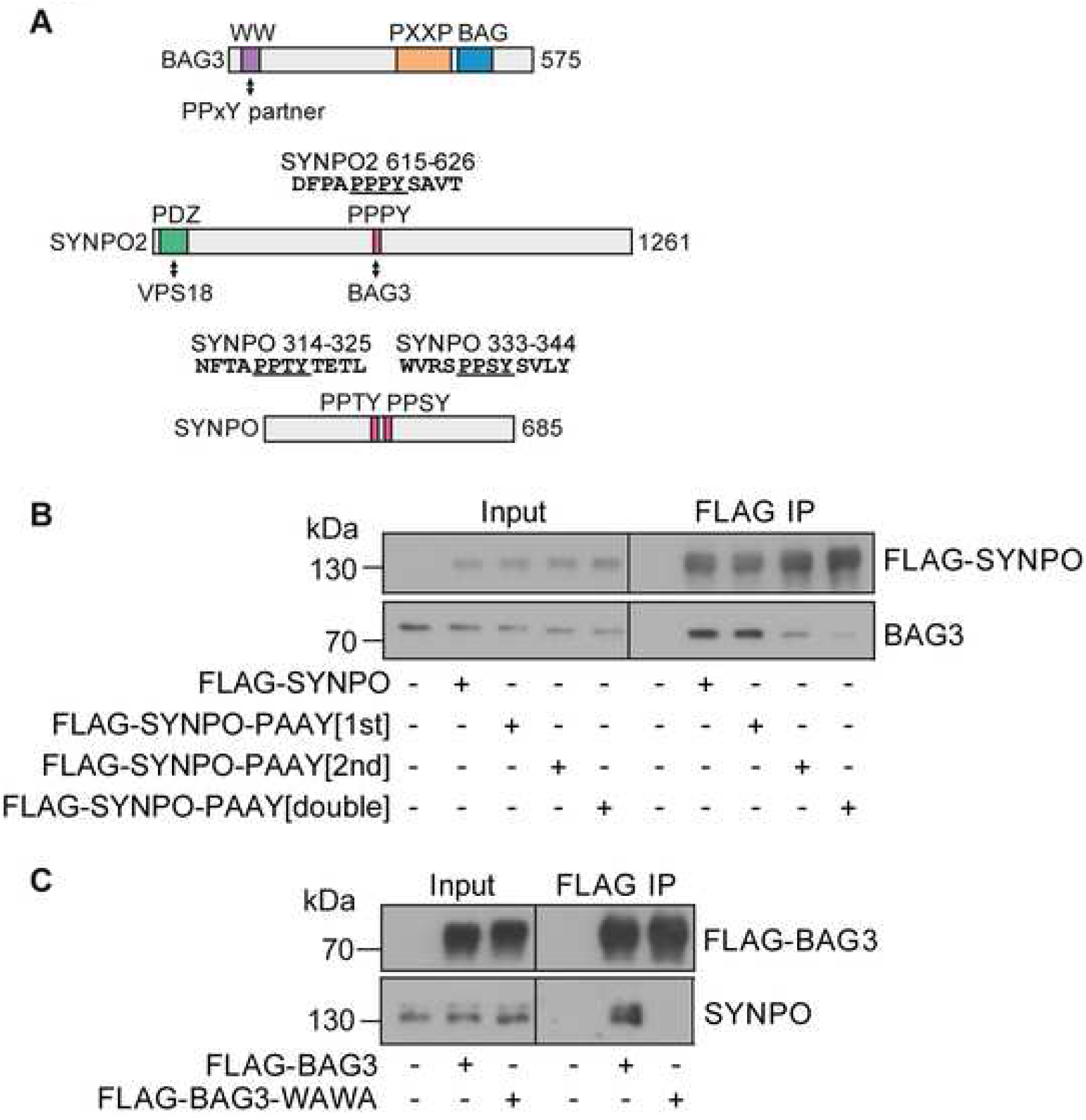
The BAG3 WW domain and SYNPO PPxY motifs are required for their interaction. (**A**) BAG3 is a multi-domain protein, which contains a WW domain at its amino-terminus for binding PPxY motifs in partner proteins. In a previously performed peptide array screen for BAG3 WW domain interacting proteins of the human proteome [18], 12-mer peptides of SYNPO2 (aa 615-626) and SYNPO (aa 333-344), respectively, were strongly recognized by the WW domain of the co-chaperone BAG3. (**B**) HeLa cells were transiently transfected with empty plasmid or plasmid constructs for the expression of FLAG-tagged SYNPO or mutant forms with inactivating mutations in the PPxY motifs, as indicated followed by immunoprecipitation with an anti-FLAG antibody (IP). Isolated immune complexes were probed for the presence of endogenous BAG3. Input samples correspond to 32 μg of protein. (**C**) Similar to the experimental approach described under (**B**), BAG3 complexes were isolated from HeLa cells expressing a wild-type form of the BAG3 co-chaperone or a form with an inactivated WW domain (BAG3-WAWA). Isolated complexes were analyzed for the presence of endogenous SYNPO.

To verify the predicted interaction, BAG3 and SYNPO, as well as mutant forms of both proteins, were expressed in HeLa cells as epitope-tagged fusion proteins and interaction was monitored by co-immunoprecipitation experiments (Figure 1B). Endogenous BAG3 was readily detectable in association with FLAG-tagged SYNPO following immunoprecipitation with an anti-FLAG antibody. A mutant form of SYNPO with the first PPTY motif inactivated (PAAY[1st]) retained the ability to interact with BAG3. In contrast, inactivation of the PPSY motif (PAAY[2nd]) strongly reduced BAG3 binding and led to complete loss of the interaction when combined with the PPTY mutation (PAAY[double]). The data demonstrate that the PPSY motif of SYNPO is the preferential BAG3 docking site, in agreement with the findings of the peptide array screen.

In a similar experimental set up, we also investigated the relevance of the BAG3 WW domain for the observed interaction. FLAG-BAG3 and FLAG-BAG3-WAWA, containing inactivating tryptophan to alanine exchanges in the WW domain, were transiently expressed in HeLa cells followed by immunoprecipitation with an anti-FLAG antibody (Figure 1C). While SYNPO was present in BAG3 complexes, it was not detectable in association with BAG3-WAWA, confirming the essential role of the WW domain of the co-chaperone in SYNPO binding. Next, we tested whether BAG3 and SYNPO interacted in primary cortical neurons. These studies revealed that endogenous SYNPO co-immunoprecipitated with BAG3 (Figure 2A). Conversely, BAG3 immunoprecipitated in the SYNPO affinity isolated fractions. Moreover, the autophagy receptor protein SQSTM1 was present in the SYNPO immunoprecipitation fractions (Figure 2B). BAG3 knockdown did not disturb the co-immunoprecipitation between SQSTM1 and SYNPO (Figure 2C). These data suggest BAG3, SYNPO and SQSTM1 are part of a functional protein complex, in which the interaction between SQSTM1 and SYNPO does not depend on the presence of BAG3.

**Figure 2.**
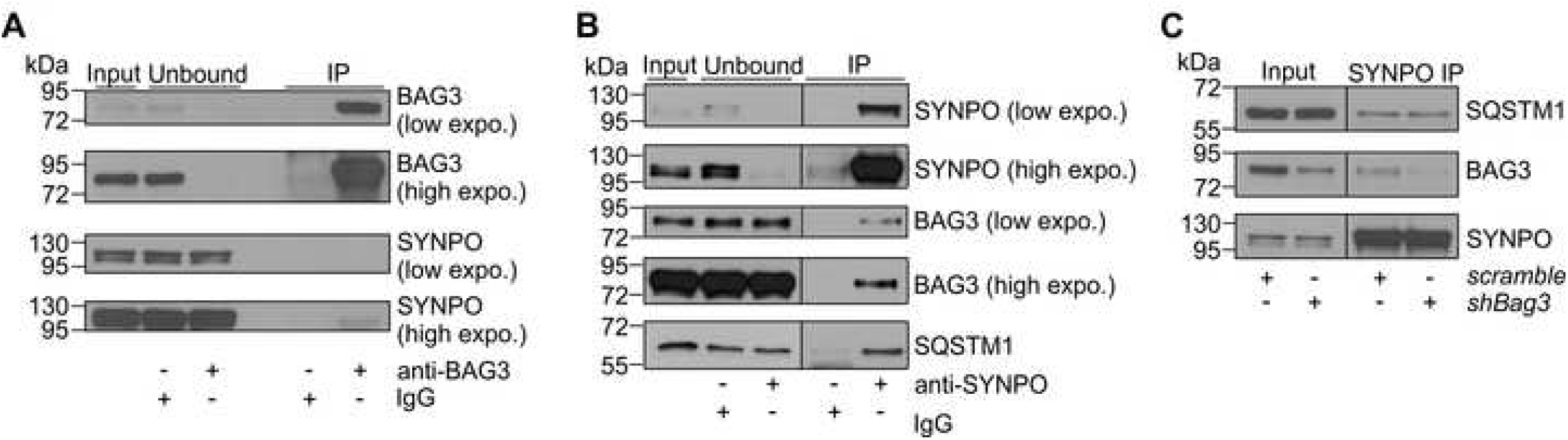
BAG3 interacts with SYNPO in mature neurons. (**A**) Immunoprecipitation of endogenous BAG3 from mature rat cortical neuronal lysates. SYNPO was detected in the isolated bound fractions. (**B**) Immunoprecipitation of endogenous SYNPO from mature rat cortical neurons. Both BAG3 and SQSTM1 were detected in the precipitated fraction. (**C**) Co-immunoprecipitation of SQSTM1 and SYNPO is independent of BAG3 in mature rat neurons.

We next investigated the spatial localization of SYNPO and BAG3 in mature primary neurons. As revealed by confocal microscopy, SYNPO immunofluorescence signals overlap with BAG3, SQSTM1 and the BAG3 interaction partner HSPA/HSP70 in neuronal processes (Figure 3A and 3C) and in the soma (Figure 3B and 3C). Colocalization analysis revealed that approximately 10% of SYNPO-positive puncta contained BAG3 (Figure 3D). Interestingly, the colocalization appeared to be in punctate-like structures (Figure 3A and 3B). Colocalization between SYNPO and MAP2 was used as a negative control (Figure 3C, 3D and Figure S1). These observations support the interaction between SYNPO and BAG3, as well as its chaperone partner HSPA/HSP70. More interestingly, SYNPO was colocalized with the autophagy receptor protein SQSTM1, suggesting the potential involvement of the BAG3-SYNPO complex in autophagy in neurons. To test this hypothesis, we further examined the spatial localization between LC3B-positive vesicles and BAG3, SYNPO or SQSTM1, respectively. Endogenous LC3B was observed in BAG3- or SYNPO-positive puncta in neuronal processes, but rarely seen in the SYN1 (synapsin I)-containing vesicles (Figure 4A and 4B). Approximately 10% of BAG3 or SYNPO were associated with LC3B-positive vesicles (Figure 4C). As expected, SQSTM1 co-stained with LC3B as well (Figure 4A-C). Taken together, these data suggest SYNPO and BAG3 are physically associated with the autophagy machinery in mature cortical neurons.

**Figure 3.**
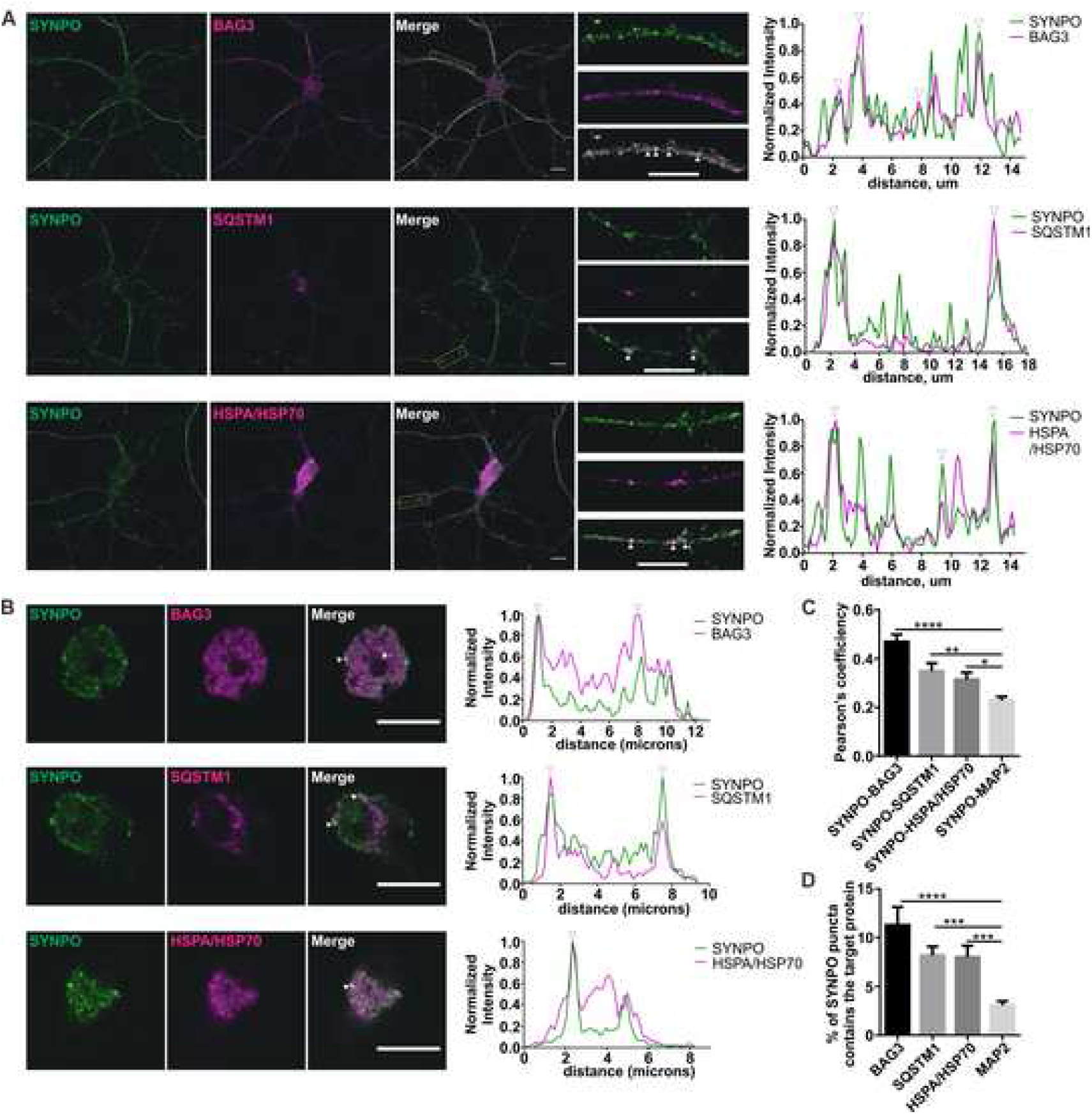
Colocalization of SYNPO with BAG3, SQSTM1 and HSPA/HSP70. Cortical neurons were immunostained for SYNPO and BAG3, SQSTM1, or HSPA/HSP70. Immunofluorescence of SYNPO overlaps with BAG3, SQSTM1 or HSPA/HSP70 in neuronal processes (**A**) and soma (**B**). Corresponding line scans are shown on the right; arrowheads indicate the areas of overlapping of intensity. Quantification of colocalization using Pearson’s correlation coefficient (**C**) and object-based analysis (**D**). In each condition, 10-30 neurons from 3 independent experiments were used for quantification. Data were plotted as mean ± SEM. As MAP2 appears in a continuous localization within neuronal dendrites and barely overlaps with SYNPO (see also Figure S1), the colocalization between SYNPO and BAG3, SQSTM1 and HSPA/HSP70, respectively, was compared to SYNPO and MAP2 using one-way ANOVA followed by Dunnett’s multiple comparisons test. ****, p < 0.0001; ***, p < 0.001; *, p < 0.05. Scale bar: 10 μm.

**Figure 4.**
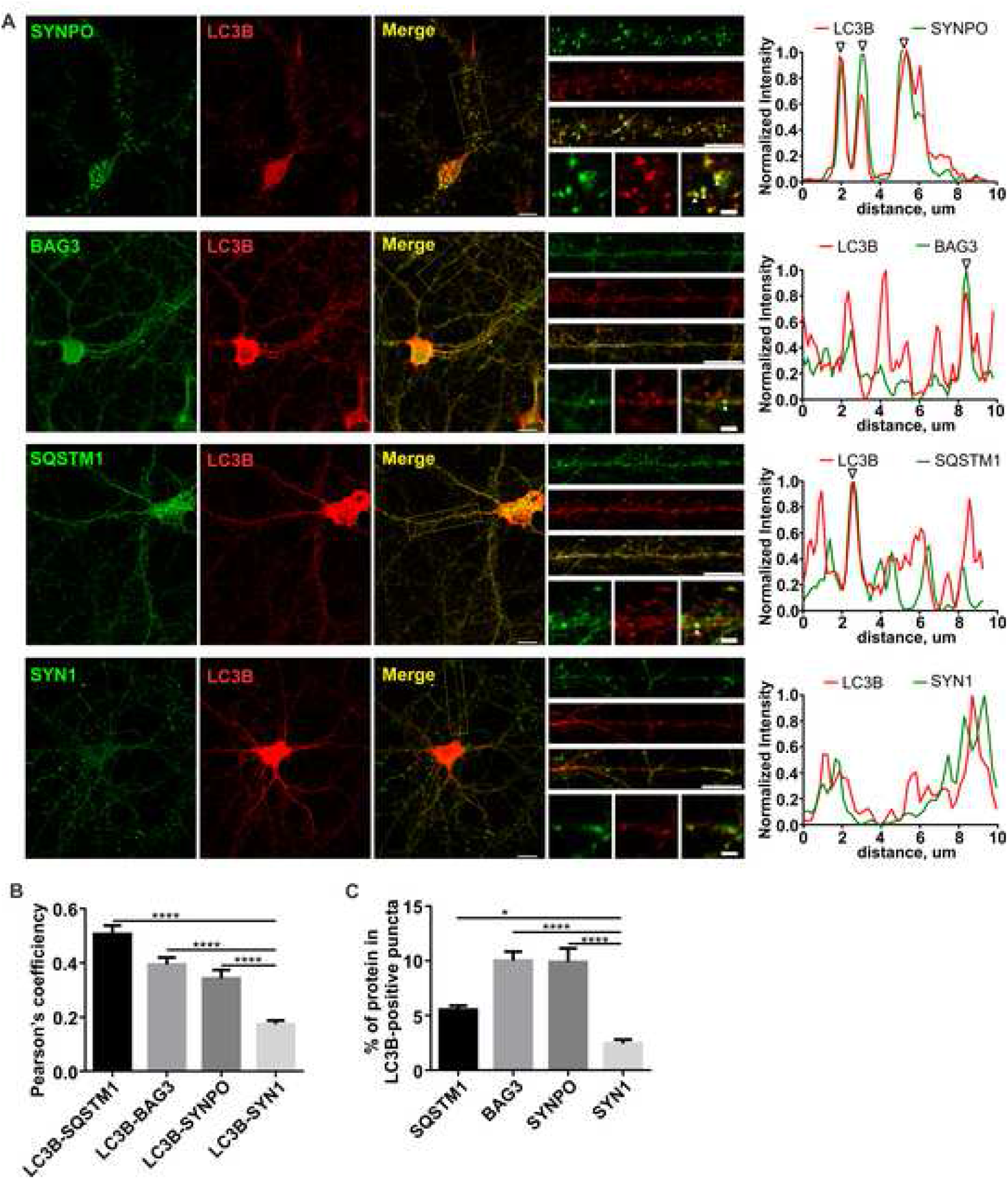
Colocalization of BAG3, SYNPO or SQSTM1 with endogenous MAP1LC3B/LC3B in neuronal processes. (**A**) Neurons were co-immunostained for LC3B and BAG3, SYNPO or SQSTM1, respectively. Overlap of BAG3, SYNPO or SQSTM1 with LC3B puncta was observed in neuronal processes. SYN1 was used as a negative control. The corresponding line scans are shown at right. Arrowheads denote areas of overlap. Scale bar: 10 μm; scale bar in the high magnification inserts: 2 μm. (**B**) Quantification of colocalization using Pearson’s correlation coefficient. (**C**) Quantification of colocalization using object based analysis. In each condition, 12-20 neurons from 3 independent experiments were used for quantification. Graphs were plotted as mean ± SEM. Colocalization between LC3B and SYNPO, BAG3 and SQSTM1, respectively, was compared to LC3B and SYN1 using one-way ANOVA followed by Dunnett’s multiple comparisons test. ****, p < 0.0001; *, p < 0.05.

### SYNPO is required for autophagosome degradation, but not autophagosome formation

The interaction between the WW domain of BAG3 and the PPPY motif in SYNPO2 is essential for autophagosome formation in muscle cells [18]. Therefore, the physical interaction of SYNPO and BAG3 prompted us to investigate if SYNPO plays a similar role in BAG3-mediated autophagy in neurons. Knockdown of either SYNPO or BAG3 increased the steady state level of LC3B-II in the absence of chloroquine, a reagent that inhibits the acidification and function of lysosomes. In contrast, knockdown of SYNPO in neurons did not result in significant changes of LC3B-II in the presence of chloroquine; however, a modest but significant decrease of LC3B-II was observed in BAG3 knockdown neurons (Figure 5A-C). Furthermore, loss of either SYNPO or BAG3 stabilized the autophagy receptor SQSTM1 (Figure 5D and 5E). Accumulation of LC3B-II and SQSTM1 suggested an inhibition of autophagosome degradation in either BAG3 or SYNPO knockdown neurons. Of note, observed changes, although statistically significant, were rather limited. This indicates that BAG3 and SYNPO dependent autophagy represents only a fraction of total autophagic flux in neurons, and might operate only at certain subcellular locations in this cell type (see below). The finding that the changes were normalized during lysosomal inhibition indicates that LC3B accumulation in the absence of BAG3 and SYNPO was not due to the enhancement of autophagy initiation, although a slight decrease of autophagy biogenesis in BAG3 knockdown was observed.

**Figure 5.**
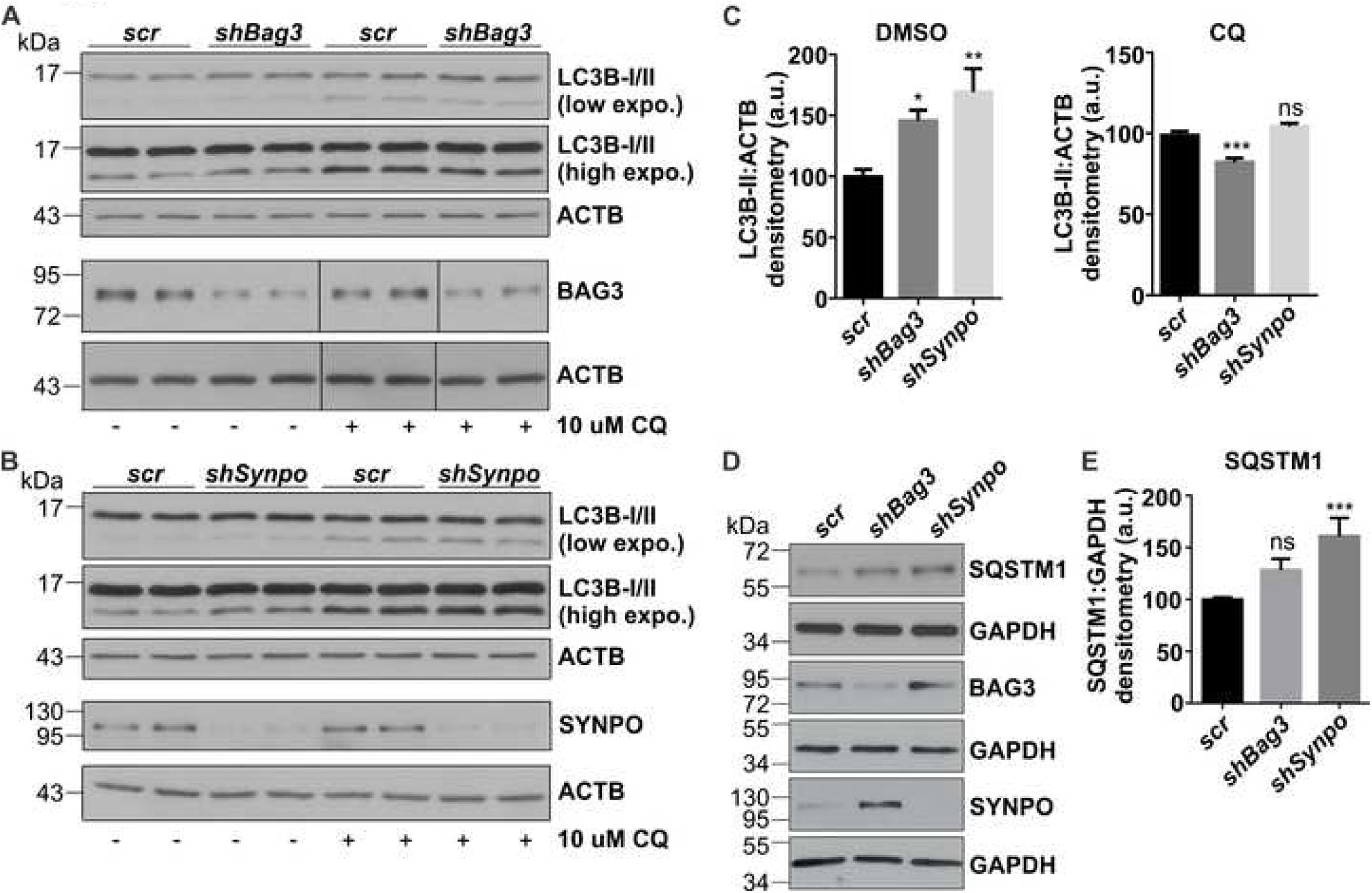
Loss of BAG3 or SYNPO reduces LC3B-II and SQSTM1 turnover. (**A**) LC3B-I/II levels in primary cortical neurons transduced with lentivirus expressing *shBag3* or a scrambled (scr) version. Neurons were treated with or without 10 μM chloroquine (CQ) for 16h. (**B**) LC3B blots of neurons transduced with lentivirus expressing shRNA for rat *Synpo* or a scrambled (*scr*) version. Neurons were treated as (**A**). (**C**) Quantifications of LC3B-II in BAG3 and SYNPO knockdown neurons in the absence or presence of CQ treatment. LC3B-II was normalized to the loading control ACTB then compared to the scrambled condition. Graph show mean ± SEM of 4-6 samples from 3 independent experiments. Statistical analysis was performed using one-way ANOVA with Tukey’s *post hoc* test. *, p<0.05; **, p<0.01; ***, p<0.001, ns, no significance. (**D**) Immunoblotting of SQSTM1, BAG3 and SYNPO in BAG3 or SYNPO knockdown neurons. GAPDH was used as loading control. (**E**) Quantification of SQSTM1 levels. Graph show mean ± SEM from 3 independent experiments. Statistical analysis was performed using one-way ANOVA with Tukey’s *post hoc* test. ***, p<0.001; ns, no significance. Scale bar: 10 μm. a.u., arbitrary units.

A block of LC3B-II degradation may be due to a decrease of autophagy flux. To test this hypothesis, we monitored autophagic flux using a tandem-fluorescence tagged reporter mTagRFP-mWasabi-LC3, which allowed us to quantify autophagosomes (green:red) and autolysosomes (red only). Knockdown of either SYNPO or BAG3 did not affected autophagic flux in neuronal soma (Figure 6A and 6C). However, autophagosomes accumulated in neuronal processes when the expression of SYNPO or BAG3 was suppressed (Figure 6B and 6C). No change was observed in autolysosomes, although the number of total autophagic vesicles significantly increased in SYNPO knockdown neurons (Figure 6B and 6C). This may indicate a compensatory induction of autophagy to counteract a block of autophagic flux at neuronal process in the absence of SYNPO, whereas such effect was not observed in BAG3 knockdown neurons. Neurons treated with bafilomycin A1 were used as positive controls. To further investigate if the blockage of autophagic flux in neuronal processes was due to an inhibition of autophagosome-lysosome fusion or lysosome dysfunction, we first examined the colocalization between LC3B and LAMP1 in neuronal processes. In control conditions, LC3B-positive vesicles were mostly colocalized with LAMP1-positive vesicles (Figure 6D and 6E). In contrast, when SYNPO expression was decreased, LC3B vesicles were proximal to LAMP1-positive lysosomes, but they did not overlap with each other (Figure 6D and E). Similarly, when the expression of BAG3 was knocked down, a decrease in overlap between LC3B and LAMP1 was observed (Figure 6D and 6E). The colocalization between GFP-LC3B and LAMP1-RFP significantly decreased in the absence of BAG3 or SYNPO (Figure 6F), suggesting that loss of BAG3 or SYNPO indeed impedes the fusion between autophagosomes and lysosomes. Next, we examined the functionality of lysosomes, which relies on the hydrolytic enzymes to become fully processed and activated at acidic pH. We tested the maturation of lysosomal protease CTSL (cathepsin L). Loss of either BAG3 or SYNPO did not change the expression pattern of CTSL compared to the control (Figure S2). Furthermore, we did not detect significant changes of the total level of LAMP1 when either SYNPO or BAG3 was depleted in neurons (data not shown). Altogether, these data suggest the interaction partners SYNPO and BAG3 are functionally involved in autophagy flux, but do not affect lysosomal function in neuronal processes.

**Figure 6.**
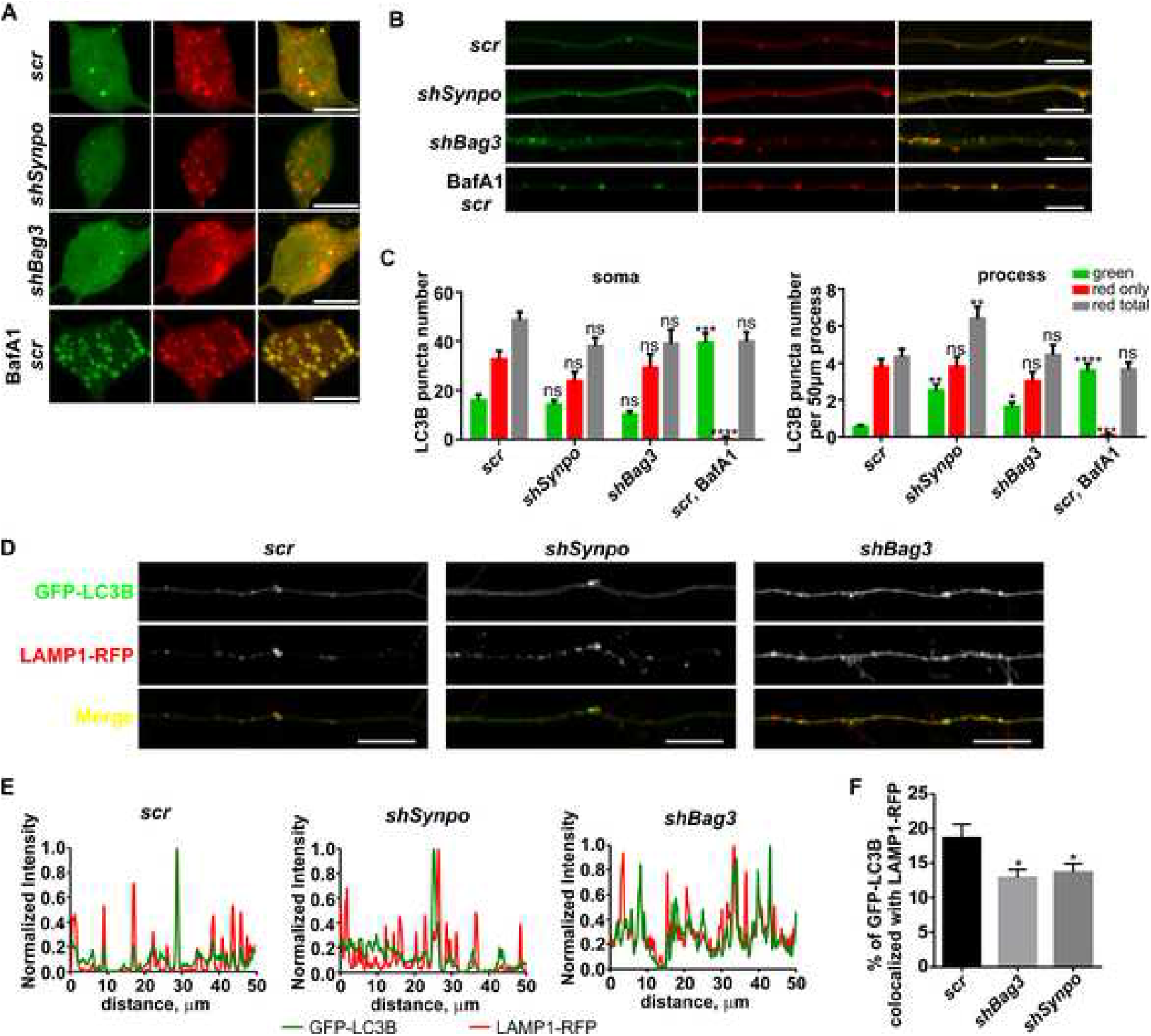
BAG3 or SYNPO knockdown blocks the autophagic flux of autophagy in neuronal processes. Representative maximal-projections of confocal z-stack images of neuronal soma (**A**) and processes (**B**). Neurons treated with 100 nM bafilomycin A1 (BafA1) for 4 h were used as positive controls. Scale bar: 10 μm. (**C**) Quantification of autophagosomes (green) and autolysosomes (red only) under the conditions of (**A**) and (**B**). The total number of green particles (autophagosomes) and red particles (autophagosomes plus autolysosomes) were counted as described in Materials and Methods. Red only particles (autolysosomes) were determined by subtracting the number of green particles from the respective number of red particles. Data were obtained from 20-30 neurons of 3 independent experiments. One to three processes from each neuron were chosen for analysis. Data were shown as mean ± SEM. Statistical analysis was performed using two-way ANOVA with Dunnett’s *post hoc* test. *, p<0.05; ****, p<0.0001; ns, no significance. (**D**) Colocalization of GFP-LC3B and LAMP1-RFP in neurons transduced with *scramble, shBag3* or *shSynpo* lentivirus. Neurons were co-transfected with GFP-LC3B and LAMP1-RFP for 48 h before fixing. Representative images are shown with corresponding line scans below (**E**). Overlap between GFP-LC3B and LAMP1-RFP decreases in either SYNPO or BAG3 knockdown neurons compared to scramble controls. Scale bar: 10 μm. (**F**) Quantification of the colocalization between GFP-LC3B and LAMP1-RFP. In each condition, data were obtained from 15-24 neurons of 2 independent experiments. 1-3 processes from each neuron were chosen for analysis. Data were shown as mean ± SEM. Statistical analysis was performed using one-way ANOVA with Dunnett’s *post hoc* test. *, p<0.05.

Interaction between SYNPO2 and BAG3 facilitates fusion of phagophore membranes in order to promote the engulfment of BAG3-client complexes [18]. SYNPO lacks a PDZ domain, which is required in SYNPO2 for contacting vacuolar protein sorting (VPS) factors in the VPS16-VPS18 membrane fusion complex [18]. To further investigate if SYNPO, in association with BAG3, also affects the maturation of autophagosomes in neurons, we performed proteinase K resistant assays and analyzed the changes of the autophagy receptor and substrate, SQSTM1 (Figure 7A). Knockdown of either SYNPO or BAG3 did not change the protection of SQSTM1 from proteinase K degradation (Figure 7B and 7C). When lysosomal degradation was inhibited by chloroquine, the levels of protected SQSTM1 were significantly increased as expected (Figure 7B and C). The extent of increase was significantly greater when BAG3 was knocked down. However, it remained the same in SYNPO knockdown, compared to the scramble control (Figure 7D). These observations suggest the autophagosomes and autolysosomes were correctly sealed in the absence of SYNPO or BAG3. Furthermore, SYNPO and BAG3 were primarily localized on the outside of autophagic vesicles, as autophagic/lysosomal inhibition did not promote the accumulation of these two proteins (Figure 7E). Taken together these data suggest that unlike SYNPO2, SYNPO in complex with BAG3 does not affect the fusion of phagophore membranes in neurons but acts at a later stage of autophagosome maturation.

**Figure 7.**
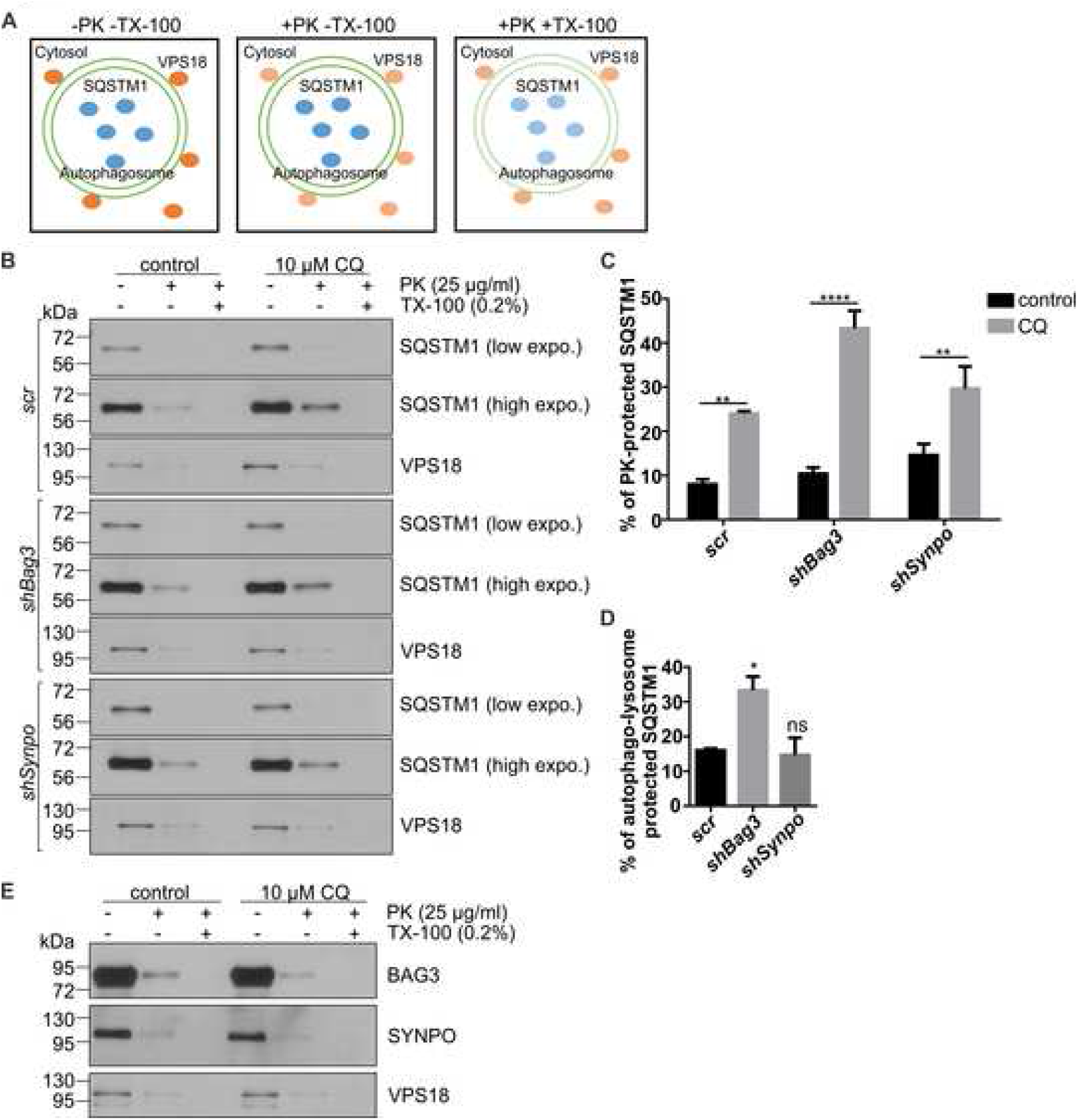
Loss of BAG3 or SYNPO does not affect the initiation or maturation of autophagosomes. (**A**) Schematic representation of proteinase K (PK) protection assay. This panel was adapted and reproduced from [46]. (**B**) Autophagic cargo receptor SQSTM1 was protected from PK digestion unless the detergent Triton X-100 (TX-100) was present. Lysates from scramble (scr), BAG3 or SYNPO knockdown neurons treated with or without 10 μM chloroquine (CQ) were subjected to PK protection assays. VPS18 was used as a cytosolic control. (**C**) Quantification of the amount of PK protected SQSTM1 in each condition. Percentage of PK protected SQSTM1 was the ratio of SQSTM1 in the presence PK but in the absence of TX-100 relative to its untreated control in a given condition. Data are shown as mean ± SEM. Statistical analysis was performed using two-way ANOVA with Tukey’s *post hoc* test. **, p<0.01; ****, p<0.0001. (**D**) Autophagosome and/or lysosome-protected SQSTM1. The difference in the amount of PK-protected SQSTM1 between chloroquine treated samples and untreated samples represent autophagosome and/or lysosome-protected SQSTM1. Data were shown as mean ± SEM. Statistical analysis was performed using one-way ANOVA with Tukey’s *post hoc* test. *, p<0.05; ns, no significance. (E) BAG3 and SYNPO were sensitive to PK digestion. Chloroquine treatment did not increase the amount of either BAG3 or SYNPO that were PK protected.

### SYNPO and BAG3 mediated autophagy facilitates the removal of phosphorylated MAPT from postsynaptic terminals

BAG3 facilitates phosphorylated MAPT autophagic degradation in young primary cortical neurons [15]. However, the underlying mechanisms remain unclear. Our data suggested that SYNPO functionally cooperates with BAG3 to support autophagic flux in mature neuronal processes at basal conditions, thus we further determined if SYNPO is also involved in MAPT degradation. Suppression of SYNPO or BAG3 expression did not change the levels of total MAPT or p-Thr231 and p-Ser396/Ser404 in mature neurons (Figure 8A and 8B). However, the levels of p-Ser262 significantly increased when either BAG3 or SYNPO was depleted (Figure 8A and 8B). The accumulation of p-Ser262 is likely due to a decrease of degradation of this MAPT species via the BAG3-SYNPO mediated autophagy. Consistently, ATG7 knockdown also resulted in increased p-Ser262 (Figure S3). Given that MAPT phosphorylated at Ser262 displays increased sorting to dendritic compartments compared to other MAPT species [28, 29], we further investigated the spatial localization of p-Ser262. This MAPT species showed increased overlap with LC3B-positive punctate in neuronal processes when either BAG3 or SYNPO was knocked down compared to the scramble control (Figure 8C and 8D). However, depletion of BAG3 or SYNPO did not increase the percentage of p-Ser262 in LC3B-positive puncta when neurons were treated with bafilomycin A1 (Figure 8C and 8D). Note that bafilomycin A1 likely affects both autophagosome-lysosome fusion and lysosomal acidification in these neurons. Altogether, these data indicate that SYNPO and BAG3 mediated autophagy facilitates the degradation of p-Ser262 in mature neuronal processes.

**Figure 8.**
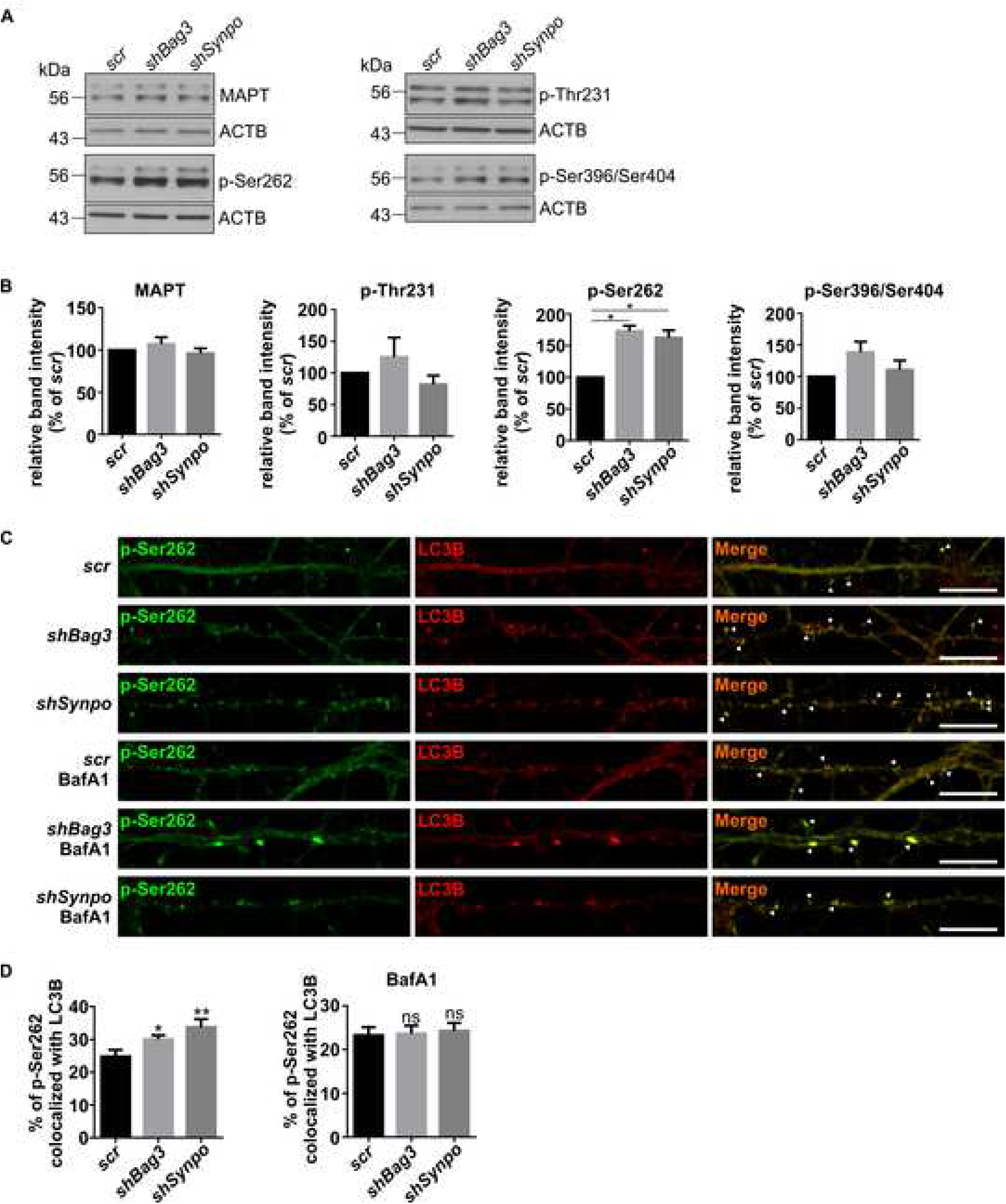
MAPT phosphorylated at Ser262 increased in neuronal processes when BAG3 or SYNPO was knocked down in mature neurons. (**A**) Representative blots of MAPT and phosphorylated MAPT (p-Thr231, p-Ser262 and p-Ser396/Ser404) in neurons transduced with *scramble (scr), shBag3* or *shSynpo* lentivirus. (**B**) Quantitation of the levels of MAPT or phosphorylated MAPT in BAG3 or SYNPO knockdown neurons from 3 independent experiments. Data were normalized to the loading control ACTB and then compared to scramble controls. Data were shown as mean ± SEM. Statistical analysis was performed using one-way ANOVA with Dunnett’s *post hoc* test. *, p<0.05. (**C**) Endogenous p-Ser262 colocalized with LC3 puncta in neuronal process in BAG3 or SYNPO knockdown neurons. Neurons were treated with DMSO or 100 nM bafilomycin A1 (BafA1) for 4 h before processing, then immunostained with anti-phosphorylated MAPT Ser262 (12E8) and LC3B antibody. Scale bar: 10 μm. Arrowheads denote the overlapping of fluorescence. (**D**) Quantification of the colocalization between phosphorylated MAPT Ser262 and LC3B. In each condition, data were obtained from 13-17 neurons of 2 independent experiments. Data were shown as mean ± SEM. Statistical analysis was performed using one-way ANOVA with Dunnett’s *post hoc* test. *, p<0.05; **, p<0.01; ns, no significance.

Given that SYNPO is a post-synaptic protein and p-Ser262 appears punctate in neuronal processes, we further investigated the subcellular localization of the accumulated p-Ser262. At basal conditions, endogenous LC3B colocalized with DLG4/PSD95 in the absence or presence of bafilomycin A1 (Figure 9A), suggesting autophagy did occur in postsynaptic compartments. When either BAG3 or SYNPO expression was suppressed, the percentage of punctate p-Ser262 within DLG4-positive compartments were significantly increased compared to the scramble controls (Figure 9B and 9C). This observation suggests p-Ser262 accumulates at postsynaptic densities when either BAG3 or SYNPO are knocked down.

**Figure 9.**
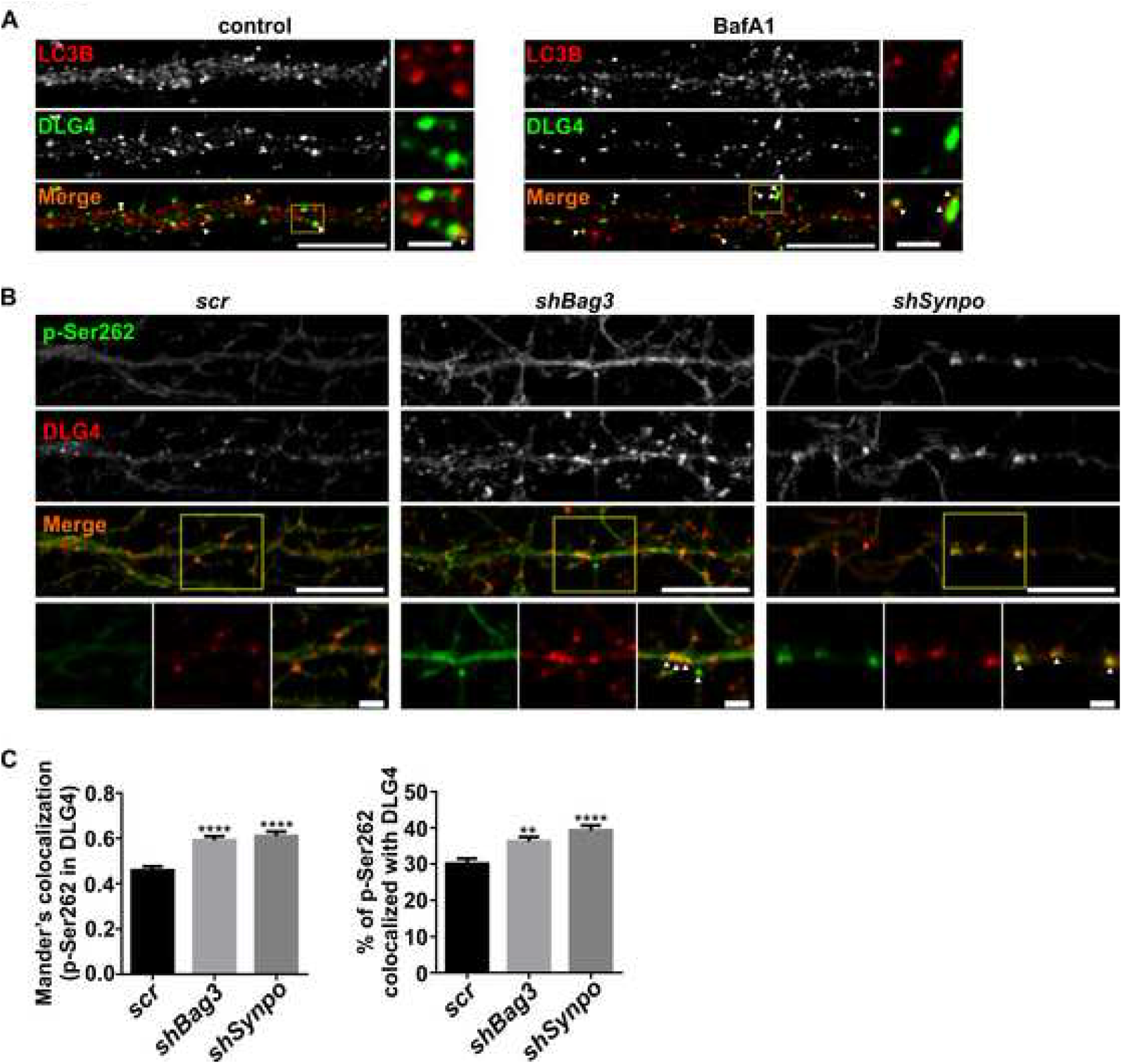
Phosphorylated MAPT Ser262 accumulates in autophagosomes at post-synaptic densities when either BAG3 or SYNPO expression is decreased. (**A**) Representative images of LC3B and DLG4/PSD95 colocalization in dendrites. Neurons were treated with either DMSO or bafilomycin A1 (BafA1) for 4 h before fixing and immunostaining. (**B**) Representative images of p-Ser262 and DLG4 co-staining in dendrites of neurons transduced with *scramble (scr), shBag3* or *shSynpo* lentivirus. Arrow heads denote the overlapping of fluorescence. (**C**) Quantification of colocalization using Mander’s colocalization coefficient and object based analysis. In each condition, 18-20 neurons from 2 independent experiments were counted. 1-3 processes from each neuron were chosen for analysis. Data were shown as mean ± SEM. Statistical analysis was performed using one-way ANOVA with Dunnett’s *post hoc* test. **, p<0.01, ****, p<0.0001. Scale bar: 10 μm. Scale bar for the high magnification insets: 2 μm.

## Discussion

Constitutive autophagy at basal levels is essential for maintaining protein homeostasis in neurons [3, 30]. Given the unique polarity of cellular structures, neuronal autophagy is regulated by compartment-specific mechanisms [5]. Efficient removal of proteins at the distal processes, as well as at dynamic synapses, is critical for neuronal health [7, 9]. In addition to previous studies focused on presynaptic autophagy [5, 8, 10], our work has revealed a regulatory mechanism of postsynaptic autophagy in facilitating the removal of phosphorylated MAPT species. Autophagic flux was supported by an interaction between the co-chaperone BAG3 and the post-synaptic scaffold protein SYNPO. Loss of either SYNPO or BAG3 resulted in accumulation of p-Ser262 in autophagic vesicles at the post-synaptic compartments. These data for the first time provide evidence of a novel role for the co-chaperone BAG3 in the post-synapse; in cooperation with SYNPO, it is part of a surveillance complex that facilitates the autophagic clearance of p-Ser262, and possibly other MAPT species. It also can be speculated that the BAG3-SYNPO complex may facilitate the clearance of other aggregation prone proteins in the dendritic compartment. It needs to be noted that our data suggest that BAG3-SYNPO-mediated autophagic processes primarily occur at the post-synaptic compartment, therefore robust global changes in phosphorylated MAPT levels and other autophagy measures (as detected by immunoblotting) would not be expected when BAG3 or SYNPO levels are manipulated. Nonetheless, the fact that significant changes are observed in autophagic processes and phosphorylated MAPT levels when BAG3 or SYNPO are knockdown demonstrate the functionality of this complex in the postsynaptic compartment.

BAG domain proteins play a vital role in protein quality control [16]. They function as cochaperones to regulate chaperone activity of HSPA/HSP70 in support of proper protein folding, preventing aggregation or targeting misfolded proteins for degradation [16]. For example, BAG1 couples HSPA/HSP70-client complexes to proteasomes for degradation [14, 16]. In contrast, BAG3 directs protein chaperone clients to autophagy for degradation [14]. As organisms age, induction of BAG3 expression in the brain likely contributes to a switch from proteasomal to autophagic degradation for the clearance of misfolded and damaged proteins [12, 31]. Clients of BAG3 include mutant huntingtin protein, misfolded superoxide dismutase 1 as well as endogenous MAPT [15, 32, 33]. All forms of MAPT species increased in immature neurons with BAG3 knockdown [15]. In our study, we observed a specific and significant accumulation of p-Ser262 when BAG3 expression was suppressed in mature neurons. This discrepancy is likely due to the fact that the mature neurons used in the present study had a significantly more extensive neuritic network and established polarity than the younger neurons used in the previous study. These data suggest that in mature neurons, BAG3 likely has a more defined role in facilitating the removal of p-Ser262 in the postsynaptic compartment.

BAG3 coordinates an array of protein partners via its modular domains to mediate the autophagic clearance of misfolded proteins [16]. The BAG domain binds with chaperone HSPA/HSP70 [16]; 2 IPV motifs directly associate with small heat shock proteins such as HSPB6 or HSPB8 [34]. Misfolded HSPA/HSP70 clients, or proteins sequestered by small heat shock proteins, are then targeted for degradation. The PXXP domain of BAG3 directly interacts with the microtubule motor dynein, which facilities retrograde trafficking and sequestration of misfolded proteins [13], and this process is regulated by YWHA/14-3-3 proteins [35]. In muscle cells, BAG3 plays a fundamental role in the autophagic degradation of mechanically unfolded and damaged cytoskeleton proteins. Through its WW domain BAG3 interacts with SYNPO2 (which contains a PDZ domain), that functionally links the BAG3 chaperone machinery to a membrane fusion complex including VPS16, VPS18 as well as STX7 (syntaxin 7). Thus a role for the BAG3-SYNPO2 interplay in the maturation of phagophore membranes was proposed [18]. We initially hypothesized a similar role for the BAG3-SYNPO interaction during neuronal autophagy. Contrary to the findings in muscle cells, loss of BAG3 in neurons did not affect the formation of the autophagosome membrane. Instead, a modest reduction of autophagosome biogenesis and a significant accumulation of autophagosomes was observed in the processes of BAG3 knockdown neurons, which is likely due in part to an inefficient fusion of autophagosomes and lysosomes. Similarly, the absence of SYNPO also did not affect the phagophore closure in mature neurons. However, the loss of SYNPO did result in a significant accumulation of autophagosomes and compensatory induction of autophagy in the mature neuronal processes. The fact that SYNPO2 has a PDZ domain and SYNPO does not, could explain why SYNPO regulates autophagy differently. Notably, BAG3 and SYNPO are multifunctional proteins, both of which participate in a variety of cellular processes; our data indicate that they functionally converge in the support of autophagosome-lysosome fusion. Consistent with this suggested role, the majority of BAG3 and SYNPO is localized on the outside of the limiting membrane of autophagosomes, and thus optimally placed to modulate membrane fusion events during autolysosome formation in the processes of mature neurons.

This study has identified SYNPO as a novel interaction partner of BAG3 that functions in concert with BAG3 to support autophagic flux in the postsynaptic compartment of mature cortical neurons. SYNPO is an actin binding protein and plays an important role in synaptic plasticity [23, 24]. The expression of SYNPO is developmentally regulated and predominantly observed in mature neurons [21]. A significant increase of SYNPO expression occurs during spine formation and stabilization [21]. SYNPO localizes at post-synaptic densities or axon initiation segments, where it contributes to the formation of the spine apparatus or cisternal organelle [22, 23, 36]. Both the spine apparatus and the cisternal organelle are extensions of the smooth endoplasmic reticulum network, which regulates local Ca^2+^ stores [26]. Physical interaction between SYNPO and BAG3 led us to explore a novel role of SYNPO in regulating autophagy in neurons.

BAG3 and SYNPO are both involved in regulation of actin-based cytoskeletal structures. Previous studies demonstrate that BAG3 plays an essential role in protein quality control [17]. In muscle cells, BAG3 senses mechanical stress and exhibits a housekeeping function by facilitating the removal of unfolded or damaged filamin [18]. During cytokinesis, BAG3 promotes the disassembly of actin-based contractile ring and actin turnover in order to assist proper cell division [37]. SYNPO also participates in the organization of the actin cytoskeleton. In neurons, formation of spine apparatus depends on SYNPO [22], which regulates Ca^2^+ release from the stores at dendritic spines [38]. How BAG3-SYNPO mediated autophagy regulates actin-based spine cytoskeleton in neurons warrants further investigations.

We observed that p-Ser262 localized to the post-synapse is a preferential target of BAG3-SYNPO-mediated autophagy in mature cortical neurons. Although MAPT is enriched in axons, it also enters dendritic spine heads in mature neurons, albeit at low levels [39, 40]. There is a growing body of literature demonstrating that dendritic MAPT plays a pivotal role in mediating signaling events at the post-synapse and inappropriate localization of MAPT to this compartment can result in neurotoxicity (for a review see [41]). For example, MAPT is required for Fyn trafficking to dendritic spines, where this kinase is essential for DLG4 recruitment by NMDA signaling [28, 39, 40]. Overexpression of frontotemporal dementia-associated P301L MAPT or MAPT pseudo-phosphorylated at T231, Ser262, or Ser396/404 promotes the trafficking of MAPT into dendritic spines in hippocampal neurons [28]. In our study, we observed a significant accumulation of the endogenous p-Ser262 in punctate structures along the neuronal processes when either SYNPO or BAG3 was suppressed, compared to the scramble controls. Among all MAPT phosphorylated species tested in this study, p-Ser262 appears to be a client that is selectively targeted by the BAG3-SYNPO complex to autophagy in mature neurons. Interestingly, the punctate p-Ser262 was closely associated with LC3B-positive vesicles as well as the post-synaptic marker DLG4. Given that the loss of SYNPO or BAG3 blocks autophagic flux in neuronal processes, the accumulation of p-Ser262 was likely due to a block of autophagy at dendritic spines in these neurons.

In conclusion, the present study further explored the mechanisms of BAG3-mediated MAPT clearance in primary neurons. SYNPO is identified as a novel interaction partner of BAG3 in neurons. The 2 proteins functionally cooperate to modulate autophagy flux and facilitate the removal of phosphorylated MAPT at dendritic spines. This may be an important pathway to regulate MAPT levels at spines in mature neurons. Given a significant reduction of SYNPO expression in pathological conditions such as Alzheimer disease and other dementias [42, 43], clearance of phosphorylated MAPT from dendritic spines may become inefficient and this may contribute to MAPT accumulation. Further studies on the pathological implications of the BAG3-SYNPO complex-mediated autophagy are needed.

## Materials and Methods

### Reagents

Constructs include: lentiviral vectors FG12-*scr* and FG12-*shBAG3* [15], viral packaging vectors psPAX2 and VSVG and pHUUG lentiviral vector for cloning (gifts from Dr. C. Pröschel, University of Rochester), GFP-LC3B (Addgene, 11546; deposited by Dr. K. Kirkegaard), mTagRFP-mWasabi-LC3 (a gift from Dr. J. Lin, Beijing University) [44], mCherry-LC3B (Addgene, 40827; deposited by Dr. D. Rubinsztein), LAMP1-RFP (Addgene, 1817; deposited by Dr. W. Mothes), pEGFP-N1 (Clontech, 6085-1). Antibodies include: rabbit anti-BAG3 (Proteintech, 10599-1-AP), goat anti-SYNPO (Santa Cruz Biotechnology, sc-21537), mouse anti-SYNPO (Santa Cruz Biotechnology, sc-515842), rabbit anti-SQSTM1 (Cell Signaling Technology, 5114) for western blotting, rabbit anti-SQSTM1 (Enzo Life Sciences, BML-PW9860) for immunofluorescence, mouse anti-GAPDH (Invitrogen, AM4300), rabbit anti-LC3B (Novus Biologicals, NB100-2220) for western blotting, rabbit anti-LC3B (Cell Signaling Technology, 2775) for immunofluorescence, mouse anti-ACTB/β-actin (Invitrogen, MA5015739), rabbit anti-VPS18 (Proteintech, 10901-1-AP), rabbit anti-MAPT (Dako, A0024), mouse anti-p-T231 (AT180; Thermo Fisher Scientific, MN1040), mouse anti-p-Ser262 (12E8, a gift from Dr. P. Dolan, Prothena Inc.), mouse anti-p-Ser396/Ser404 (PHF1, a gift from Dr. P. Davies), rabbit anti-HSP70 (Assay Designs, SPA-757), rabbit anti-DLG4/PSD95 (Cell Signaling Technology, 3450), mouse anti-DLG4/PSD95 (Santa Cruz Biotechnology, sc-32290), mouse anti-FLAG (clone M2; Sigma-Aldrich, F1804) and M2-agarose (Sigma-Aldrich, A1205), rabbit anti-SYN1 (D12G5; Cell Signaling Technology, 5297), rabbit anti-MAP2 (a gift from Dr. H.Fu, Ohio State University).

### Cell culture

Primary cortical neurons: Timed pregnant Sprague-Dawley female rats were purchased from Charles River laboratories. Animals were housed and handled according to the protocols approved by the Institutional Animal Care and Use Committee (IACUC) at the University of Rochester. Primary cortical neurons were prepared as previously described [45]. Briefly, cortices from embryonic day 18 pups were pooled together and dissociated in 0.25% trypsin for 20 min at 37°C. Following gently trituration, neurons were plated at a high density of 100,000 cells/cm^2^ for biochemical studies, and at a lower density of 15,000 cells/cm^2^ for imaging. Neurons were grown for 20-24 days in vitro (DIV) in maintenance medium (Neurobasal medium (Thermo Fisher Scientific 21103049) supplemented with 2% B27 (Thermo Fisher Scientific 17504044) and 2 mM GlutaMax (Thermo Fisher Scientific 35050061). To inhibit glia proliferation, 1.25 μM cytosine arabinoside (AraC; Sigma Aldrich, C1768) was added at DIV4; thereafter, 30% of media was replaced every 3-4 days. For lentiviral transduction, DIV16 neurons were treated with virus in a half volume of growth media for 16-24 h; then, the conditioned medium supplemented with an equal volume of fresh medium was added back. For transfection, neurons grown on #1.5 18-mm coverslips was transfected with 1.5 μg DNA using Lipofectamine 2000 (Thermo Fisher Scientific, 11668027) for 4-6 h, then fresh media was added back for other 36-48 h.

HeLa cell culture: HeLa cells were grown in DMEM medium supplemented with 10% fetal bovine serum, 2 mM glutamine and 100 U penicillin/ 0.1 mg/ml streptomycin on 6 cm dishes until 70% confluent. HeLa cells were transfected with 3 μg of the designated construct or empty plasmid (see below) using the CalPhos transfection kit (Clontech, 631312). HeLa cells were used 48 h after transfection.

### Lentivirus

Short-hairpin RNA containing siRNA (5’-GGAAATCAATGTTCACGTTCG-3’) that targets all isoforms of rat *Synpo* or siRNA (5’-GCAAGCGAAAGCTGGTCATCA-3’) targeting rat *Atg7* was cloned into the pHUUG vector. To generate lentiviral particles, lentiviral vectors were cotransfected with viral packaging vectors into 70% confluent HEK293TN cells using PolyJet (SignaGen Laboratories, SL100688). Media were changed into DMEM containing 1% FetalClone-II serum (Thermo Fisher Scientific, SH3006603) 16 h after transfection. The transfected HEK293TN cells were then incubated at 33°C and 5% CO2 to slow their proliferation. Virus was collected at 64 h after transfection and filtered through a 0.2 μm syringe filter to remove cell debris. Viral media were concentrated by ultracentrifugation at 4°C, and resuspended in Neurobasal medium by gently pipetting. Concentrated viral stocks were aliquoted, snap frozen and stored at −80°C.

### Immunoprecipitation

Neurons grown on 6 cm dishes were lysed in ice-cold lysis buffer (50 mM Tris, 150 mM NaCl, 0.4% NP-40 [Sigma Aldrich, 492016] 1 mM EDTA, 1 mM EGTA, pH 7.4) supplemented with protease inhibitor cocktail (Sigma Aldrich, P8340) and phosphatase inhibitors (Sigma Aldrich, P0044). Cell lysates were briefly sonicated then centrifuged at 15,000 x g for 10 min at 4°C. Cleared supernatants (500 μg) were mixed with 2 μg IgG control (Sigma Aldrich, 12-371 (normal mouse IgG control) or 12-370 (normal rabbit IgG control)) or primary antibody, followed by incubation at 4°C for 24 h. The next day, antibody/antigen mixture was incubated with Dynabeads M-280 sheep anti-rabbit IgG (Thermo Fisher Scientific, 11203D) or Dynabeads M-280 sheep anti-mouse IgG (Thermo Fisher Scientific, 11201D) for 6 h at 4°C. An aliquot of unbound proteins was saved. After thoroughly washing the beads, bound fractions were eluted in sample loading buffer by boiling at 100°C for 10 min. Proteins were then resolved by SDS-PAGE.

To investigate the contribution of the PPTY and PPSY motif of SYNPO to BAG3 binding, the motifs were changed to PAAY separately and in combination (see Figure 1) by PCR-based site directed mutagenesis using the human muscle isoform of *SYNPO* as template (NP_001159681), followed by subcloning into plasmid pCMV2b (Agilent Technologies, 211172) for expression of FLAG-tagged fusion proteins. Transfected cells were lysed in 25 mM MOPS (Omnipur, 6310), pH 7.2, 100 mM KCl and 0.5 % Tween 20 (Amresco, 5000183) (buffer A) containing 5 mM EDTA and complete protease inhibitors (Roche, 11836153001). The lysate was centrifuged at 30,000 x g for 20 min at 4°C and the soluble fraction was used for immunoprecipitation. Samples were incubated with M2-agarose (Sigma-Aldrich, A2220) at a concentration of 30 μl of agarose per ml of lysate for 3 h at 4°C. The agarose was collected by centrifugation and washed 5 times with buffer A and once with buffer A lacking detergent. Agarose-associated proteins were eluted with 0.1 M glycine, pH 3.5, diluted with sample loading buffer and incubated at 100°C for 10 min. prior to SDS-PAGE. To verify the role of the BAG3 WW domain in SYNPO binding, human BAG3 and BAG3-WAWA (carrying tryptophan to alanine substitutions at position 26 and 49) were expressed in HeLa cells as FLAG-tagged versions, followed by immunoprecipitation of co-chaperone complexes as described above.

### Proteinase K resistance assay

Neurons were grown on 6 cm dishes and treated with or without 10 μM chloroquine (Fisher Scientific, ICN19391980) in maintenance medium for 16 h. The protocol for the proteinase protection assay was adapted from a previous study [46]. Briefly, neurons were homogenized in ice-cold detergent-free buffer (20 mM HEPES, 220 mM mannitol, 70 mM sucrose, 1 mM EDTA, pH 7.6) after treatment. After centrifugation at 500 x g for 5 min, the post-nuclear supernatant was collected and divided equally into 3 clean microfuge tubes. One tube was left untreated, while the other 2 were treated with 25 μg/ml proteinase K (Sigma Aldrich, P2308) in the absence or presence of 0.2% (v:v) Triton X-100 (Fisher Scientific, BP151-100) for 10 min on ice. Proteinase K was then stopped by adding protease inhibitors (1 mM PMSF, 10 μg/ml of aprotinin, leupeptin and pepstatin [Fisher Scientific, 44-865-0, AAJ11388MB, 10-897-625MG and 51-648-125MG, respectively]). All samples were precipitated with 10% trichloracetic acid. Protein pellets were then resuspended in SDS sample buffer and prepared for immunoblotting.

### Immunoblotting

Depending on the antibody to be used, 20 to 40 μg total protein was denatured in 1x SDS sample buffer at 100°C for 10 min before being loaded onto each lane of 10% SDS-PAGE gels. After electrophoresis, proteins were transferred onto nitrocellulose membranes. Membranes were then blocked in TBST (20 mM Tris, 137 mM NaCl, pH 7.6 0.1% Tween-20) containing 5% non-fat dry milk for 1 h at room temperature. Primary antibodies were diluted in blocking solutions as follows: rabbit anti-BAG3, 1:3000; mouse anti-FLAG, 1:1000; goat anti-SYNPO, 1:200; mouse anti-SYNPO, 1:1000; rabbit anti-SQSTM1, 1:1000; mouse anti-GAPDH, 1:10000; rabbit anti-LC3B, 1:5000; mouse anti-ACTB, 1:10000; rabbit anti-VPS18, 1:1000; rabbit anti-MAPT, 1:10,000; mouse anti-p-Thr231 (AT180), 1:1000; mouse anti-p-Ser262 (12E8), 1:1000; mouse anti-p-Ser396/Ser404 (PHF1), 1:5000, followed by incubation at 4°C overnight. The next day, membranes were further incubated with secondary antibody for 1 h at room temperature. After thoroughly washing, membranes were visualized by enhanced chemiluminescence. The intensity of each band was quantified using Image Studio Lite (Li-Cor). GAPDH or ACTB was used as loading controls. Treatments were then normalized to their corresponding control sample and expressed as a fold difference above the control.

### Immunofluorescence

Neurons grown on coverslips were rinsed with phosphate buffered saline (PBS), followed by fixing in PBS containing 4% paraformaldehyde and 4% sucrose for 10 min at room temperature. Then, cells were permeabilized in PBS containing 0.25% Triton X-100 for another 10 min at room temperature. Thereafter, non-specific binding sites were blocked with PBS containing 5% BSA (Fisher Scientific, AAJ1085718) and 0.3 M glycine. Primary antibodies were diluted in blocking solution as follows: rabbit anti-BAG3, 1:500; goat anti-SYNPO, 1:100; mouse anti-SYNPO, 1:100; rabbit anti-SQSTM1, 1:500; rabbit anti-HSP70, 1:200; rabbit anti-DLG4/PSD95; mouse anti-DLG4/PSD95, 1:100; mouse anti-p-Ser262 (12E8), 1:100; rabbit anti-MAP2, 1:1000; rabbit anti-SYN1, 1:250, followed by incubation at 4°C overnight. The next day, neurons were washed with PBS 3 times. Then, Alexa Fluor 488-conjugated donkey anti-mouse IgG (Invitrogen, A-21202) or Alexa Fluor 594-conjugated donkey anti-rabbit IgG (Invitrogen, A-21207) was diluted in blocking solution and incubated with neurons for 1 h at room temperature. After thoroughly washing, coverslips were mounted onto ProLong Diamond Antifade Mountant (Thermo Fisher Scientific, P36965). Images were acquired on an Olympus laser scanning confocal microscope using a 60x 1.35 numerical aperture oil-immersion objective with a 1.5x optical magnification.

### mTagRFP-mWasabi-LC3 reporter assay

Neurons grown on coverslips were transfected with mTagRFP-mWasabi-LC3 plasmid for 36-48 h using Lipofectamine 2000 (Thermo Fisher Scientific, 11668019). Transfected neurons were washed with warm PBS once before being transferred into imaging buffer (137 mM NaCl, 2.7 mM KCl, 10 mM Na2HPO4, 1.8 mM KH2PO4, 1 mM CaCl2, 0.5 mM MgCl2, 5 mM glucose, pH 7.4). Images were acquired on an Olympus laser scanning confocal microscope with a 40x 1.35 numerical aperture oil-immersed objective with a 2.0x optical magnification. A stack of images was acquired with a z step interval of 0.2 μm.

### Image analysis

Images were opened and processed in Fiji (https://fiji.sc/). Background was subtracted by the rolling ball method in Fiji with a radius of 50 μm. Region of interests were either selected around soma, or a line (line width equals 75 pixels) was drawn along processes followed by straightening. Line scans were generated using the plot profile tool in Fiji (line width of one pixel). Intensity values for a given marker were normalized by subtracting the minimum, and dividing the dataset by its maximum. To measure the total number of mWasabi-LC3 or mTagRFP-LC3, maximum projections were generated in Fiji. LC3B-positive area was thresholded and segmented using watershed algorithm in Fiji. The total number of LC3B particles was measured using the analyze particles function. Colocalization analyses were performed using JAcoP plugin in Fiji [47, 48]. Specifically, background was subtracted from each channel using the roll ball method with a radius of 50 μm. Regions of interest were selected as the entire image (Figure 3 and 4) or along processes with a line width of 75 pixels followed by straightening (Figure 6, 8 and 9). Immuno-positive area was then thresholded using the build-in function in JAcoP. Both intensity based (Pearson’s correlation coefficient and Mander’s colocalization coefficients) and object based methods (centers of mass-particles coincidence) were used to quantify the overlapping of fluorescence intensity. Pearson’s correlation coefficient and Mander’s colocalization coefficients were calculated in JAcoP with the default settings. Centers of mass-particles coincidence was measured in JAcoP with a limitation of particle size between 3-200 pixels.

### Statistical analysis

All image measurements were obtained from the raw data. GraphPad Prism was used to plot graphs and perform statistical analysis. The statistical tests were denoted in each figure legend. Images were prepared and assembled in CorelDraw graphic suite. Brightness and contrast were adjusted equally for all images within a panel.

## Acknowledgements

The authors would also like to thank Dr. D. Yule for the use of his confocal microscope. Dr. C. Pröschel is thanked for providing us with the lentiviral vectors and Dr. Lin for providing us with the mTagRFP-mWasabi-LC3 vector. We thank Dr. P. Davies for the PHF1 antibody, Dr. P. Dolan for the 12E8 antibody and Dr. Fu for the MAP2 antibody.

---

**Figure S1.**
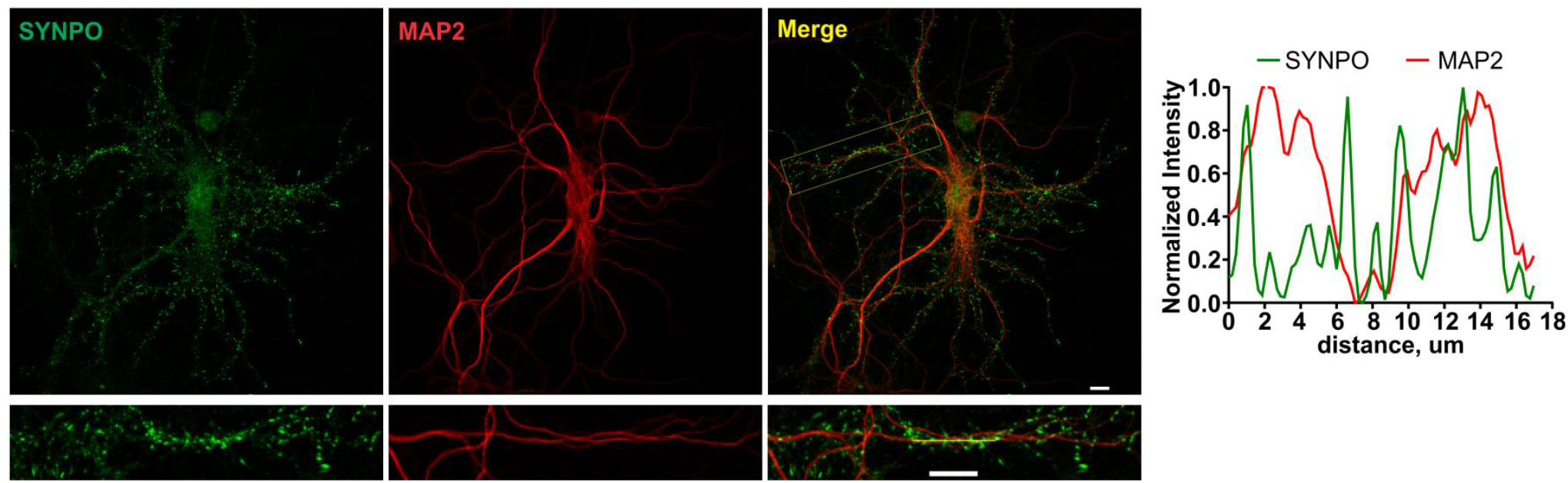
SYNPO does not show significant overlap with MAP2. Immunofluorescence images of mature neurons stained for SYNPO (green) and MAP2 (red). Merged images shows that SYNPO shows little overlap with MAP2 in neuronal processes (images in the bottom row). Corresponding line scanning of a process is shown at the right. Scale bar: 10 μm.

**Figure S2.**
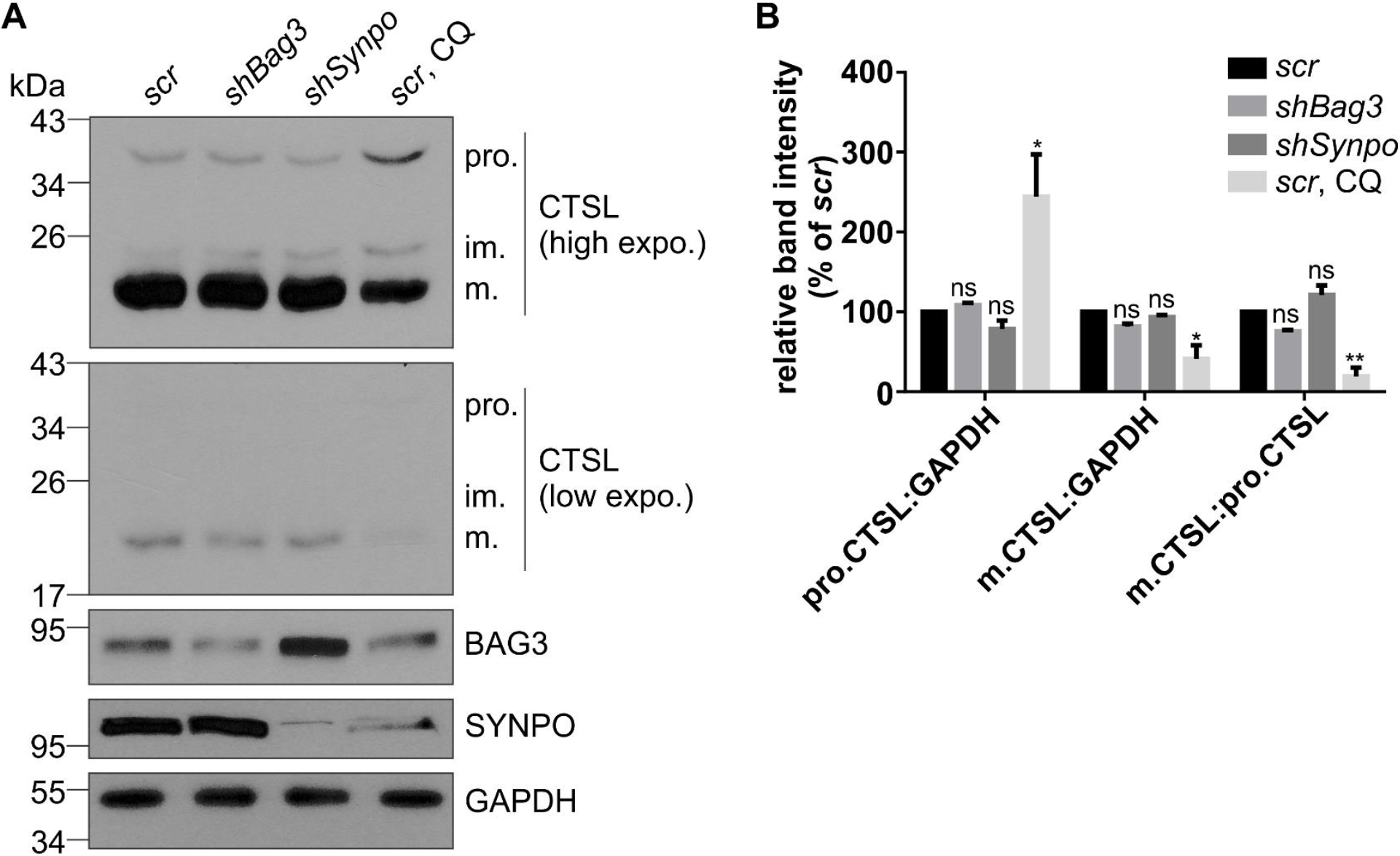
SYNPO or BAG3 knockdown does not affect lysosomal function. (**A**) Maturation of CTSL (cathepsin L) in either BAG3 or SYNPO knockdown neurons by immunoblotting. pro., precursor CTSL; im, immature CTSL; m, mature CTSL. Neurons treated with 10 μM chloroquine (CQ) were used as a positive control. (**B**) Quantification of precursor CTSL:mature CTSL ratio. Graph shows mean ± SEM. Statistical analysis was performed using one-way ANOVA with Dunnett’s *post hoc* test. **, p<0.01; *, p<0.05; ns, no significance.

**Figure S3.**
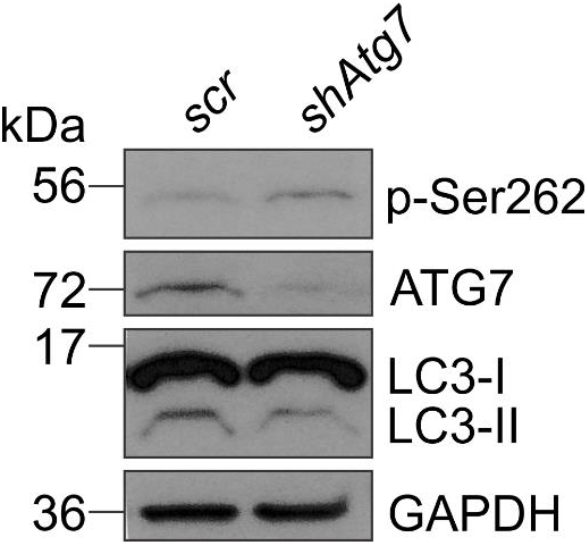
Immunoblot of MAPT phosphorylated at Ser262 (12E8) in ATG7 knockdown neurons. Mature rat cortical neurons were transduced with lentivirus expression *shAtg7* or *scrambled* shRNA. The level of p-Ser262 increased, while LC3B-II decreased in ATG7 knockdown neurons compared to the control. GAPDH was used as a loading control.

## Abbreviations

aa: amino acids
ACTB: β actin
BafAl: bafilomycin Aι
BAG3: BCL2 associated athanogene 3
CQ: chloroquine
CTSL: cathepsin L
DIV: days in vitro
DLG4/PSD95: discs large MAGUK scaffold protein 4
HSPA/HSP70: heat shock protein family A (Hsp70)
MAP1LC3B/LC3B: microtubule associated protein 1 light chain 3 beta
MAP2: microtubule associated protein 2
MAPT: microtubule associated protein tau
p-Ser262: MAPT phosphorylated at serine 262
p-Ser396/404: MAPT phosphorylated at serines 396 and 404
p-Thr231: MAPT phosphorylated at threonine 231
PBS: phosphate buffered saline
PK: proteinase K
scr: scrambled
shRNA: short hairpin RNA
SQSTM1/p62: sequestosome 1
SYN1: synapsin 1
SYNPO: synaptopodin
SYNPO2: synaptopodin-2 (myopodin)
VPS: vacuolar protein sorting

## References

1. Ariosa, A.R. and D.J. Klionsky, Autophagy core machinery: overcoming spatial barriers in neurons. J Mol Med (Berl), 2016. 94(11): p. 1217–1227.

2. Galluzzi, L., et al., Molecular definitions of autophagy and related processes. Embo J, 2017. 36(13): p. 1811–1836.

3. Hara, T., et al., Suppression of basal autophagy in neural cells causes neurodegenerative disease in mice. Nature, 2006. 441(7095): p. 885–9.

4. Komatsu, M., et al., Loss of autophagy in the central nervous system causes neurodegeneration in mice. Nature, 2006. 441(7095): p. 880–4.

5. Maday, S. and E.L. Holzbaur, Compartment-specific regulation of autophagy in primary neurons. J Neurosci, 2016. 36(22): p. 5933–45.

6. Fu, M.M. and E.L. Holzbaur, MAPK8IP1/JIP1 regulates the trafficking of autophagosomes in neurons. Autophagy, 2014. 10(11): p. 2079–81.

7. Soukup, S.-F., et al., A LRRK2-dependent endophilinA phosphoswitch is critical for macroautophagy at presynaptic terminals. Neuron, 2016. 92(4): p. 829–844.

8. Maday, S. and E.L. Holzbaur, Autophagosome biogenesis in primary neurons follows an ordered and spatially regulated pathway. Dev Cell, 2014. 30(1): p. 71–85.

9. Vanhauwaert, R., et al., The SAC1 domain in synaptojanin is required for autophagosome maturation at presynaptic terminals. Embo J, 2017. 36(10): p. 1392-1411.

10. Okerlund, N.D., et al., Bassoon controls presynaptic autophagy through Atg5. Neuron, 2017. 93(4): p. 897–913.e7.

11. Stavoe, A.K., et al., KIF1A/UNC-104 transports ATG-9 to regulate neurodevelopment and autophagy at synapses. Dev Cell, 2016. 38(2): p. 171–85.

12. Gamerdinger, M., et al., Protein quality control during aging involves recruitment of the macroautophagy pathway by BAG3. Embo J, 2009. 28(7): p. 889–901.

13. Gamerdinger, M., et al., BAG3 mediates chaperone-based aggresome-targeting and selective autophagy of misfolded proteins. EMBO Rep, 2011. 12(2): p. 149–56.

14. Minoia, M., et al., BAG3 induces the sequestration of proteasomal clients into cytoplasmic puncta: implications for a proteasome-to-autophagy switch. Autophagy, 2014. 10(9): p. 1603–21.

15. Lei, Z., C. Brizzee, and G.V.W. Johnson, BAG3 facilitates the clearance of endogenous Tau in primary neurons. Neurobiol Aging, 2015. 36(1): p. 241–248.

16. Behl, C., Breaking BAG: the co-chaperone BAG3 in health and disease. Trends Pharmacol Sci, 2016. 37(8): p. 672–88.

17. Klimek, C., et al., BAG3-mediated proteostasis at a glance. J Cell Sci, 2017. 130(17): p. 2781–2788.

18. Ulbricht, A., et al., Cellular mechanotransduction relies on tension-induced and chaperone-assisted autophagy. Curr Biol, 2013. 23(5): p. 430–5.

19. Merabova, N., et al., WW domain of BAG3 is required for the induction of autophagy in glioma cells. J Cell Physiol, 2015. 230(4): p. 831–41.

20. Mundel, P., et al., Synaptopodin: an actin-associated protein in telencephalic dendrites and renal podocytes. J Cell Biol, 1997. 139(1): p. 193–204.

21. Czarnecki, K., et al., Postnatal development of synaptopodin expression in the rodent hippocampus. J Comp Neurol, 2005. 490(2): p. 133–44.

22. Deller, T., et al., Synaptopodin-deficient mice lack a spine apparatus and show deficits in synaptic plasticity. Proc Natl Acad Sci U S A, 2003. 100(18): p. 10494–9.

23. Kremerskothen, J., et al., Synaptopodin, a molecule involved in the formation of the dendritic spine apparatus, is a dual actin/alpha-actinin binding protein. J Neurochem, 2005. 92(3): p. 597–606.

24. Vlachos, A., et al., Synaptopodin regulates plasticity of dendritic spines in hippocampal neurons. J Neurosci, 2009. 29(4): p. 1017–33.

25. Asanuma, K., et al., Synaptopodin orchestrates actin organization and cell motility via regulation of RhoA signalling. Nat Cell Biol, 2006. 8(5): p. 485–91.

26. Korkotian, E. and M. Segal, Synaptopodin regulates release of calcium from stores in dendritic spines of cultured hippocampal neurons. J Physiol, 2011. 589(Pt 24): p. 5987–95.

27. Iwasaki, M., et al., BAG3 directly associates with guanine nucleotide exchange factor of Rap1, PDZGEF2, and regulates cell adhesion. Biochem Biophys Res Commun, 2010. 400(3): p. 413–8.

28. Xia, D., C. Li, and J. Gotz, Pseudophosphorylation of Tau at distinct epitopes or the presence of the P301L mutation targets the microtubule-associated protein Tau to dendritic spines. Biochim Biophys Acta, 2015. 1852(5): p. 913–24.

29. Zempel, H., et al., Abeta oligomers cause localized Ca(2+) elevation, missorting of endogenous Tau into dendrites, Tau phosphorylation, and destruction of microtubules and spines. J Neurosci, 2010. 30(36): p. 11938–50.

30. Boland, B., et al., Autophagy induction and autophagosome clearance in neurons: relationship to autophagic pathology in Alzheimer’s disease. J Neurosci, 2008. 28(27): p. 6926–37.

31. Tang, M., et al., Nrf2 mediates the expression of BAG3 and autophagy cargo adaptor proteins and Tau clearance in an age-dependent manner. Neurobiol Aging, 2017. 63: p. 128–139.

32. Carra, S., et al., HspB8 chaperone activity toward poly(Q)-containing proteins depends on its association with Bag3, a stimulator of macroautophagy. J Biol Chem, 2008. 283(3): p. 1437–44.

33. Crippa, V., et al., The small heat shock protein B8 (HspB8) promotes autophagic removal of misfolded proteins involved in amyotrophic lateral sclerosis (ALS). Hum Mol Genet, 2010.19(17): p. 3440–56.

34. Rauch, J.N., et al., BAG3 is a modular, scaffolding protein that physically links heat shock protein 70 (HSP70) to the small heat shock proteins. J Mol Biol, 2017. 429(1): p. 128–141.

35. Xu, Z., et al., 14-3-3 protein targets misfolded chaperone-associated proteins to aggresomes. J Cell Sci, 2013. 126(Pt 18): p. 4173–86.

36. Schluter, A., et al., Structural plasticity of synaptopodin in the axon initial segment during visual cortex development. Cereb Cortex, 2017. 27(9): p. 4662–4675.

37. Varlet, A.A., et al., Fine-tuning of actin dynamics by the HSPB8-BAG3 chaperone complex facilitates cytokinesis and contributes to its impact on cell division. Cell Stress Chaperones, 2017. 22(4): p. 553–567.

38. Korkotian, E., M. Frotscher, and M. Segal, Synaptopodin regulates spine plasticity: mediation by calcium stores. J Neurosci, 2014. 34(35): p. 11641–51.

39. Hoover, B.R., et al., Tau mislocalization to dendritic spines mediates synaptic dysfunction independently of neurodegeneration. Neuron, 2010. 68(6): p. 1067–81.

40. Ittner, L.M., et al., Dendritic function of Tau mediates amyloid-beta toxicity in Alzheimer’s disease mouse models. Cell, 2010. 142(3): p. 387–97.

41. Ittner, A. and L.M. Ittner, Dendritic Tau in Alzheimer’s Disease. Neuron, 2018. 99(1): p. 13–27.

42. Counts, S.E., et al., Synaptic gene dysregulation within hippocampal CA1 pyramidal neurons in mild cognitive impairment. Neuropharmacology, 2014. 79: p. 172–9.

43. Datta, A., et al., An iTRAQ-based proteomic analysis reveals dysregulation of neocortical synaptopodin in Lewy body dementias. Mol Brain, 2017. 10(1): p. 36.

44. Zhou, C., et al., Monitoring autophagic flux by an improved tandem fluorescent-tagged LC3 (mTagRFP-mWasabi-LC3) reveals that high-dose rapamycin impairs autophagic flux in cancer cells. Autophagy, 2012. 8(8): p. 1215–26.

45. Ji, C., M. Tang, and G.V.W. Johnson, Assessing the degradation of Tau in primary neurons: The role of autophagy. Methods Cell Biol, 2017. 141: p. 229–244.

46. Nguyen, T.N., et al., Atg8 family LC3/GABARAP proteins are crucial for autophagosome-lysosome fusion but not autophagosome formation during PINK1/Parkin mitophagy and starvation. The Journal of Cell Biology, 2016. 215(6): p. 857–874.

47. Dunn, K.W., M.M. Kamocka, and J.H. McDonald, A practical guide to evaluating colocalization in biological microscopy. Am J Physiol Cell Physiol, 2011. 300(4): p. C723–42.

48. Bolte, S. and F.P. Cordelieres, A guided tour into subcellular colocalization analysis in light microscopy. J Microsc, 2006. 224(Pt 3): p. 213–32.

